# A computational approach for deciphering the interactions between proximal and distal regulators in B cell differentiation

**DOI:** 10.1101/2023.11.02.565268

**Authors:** Sung-Joon Park, Kenta Nakai

**Affiliations:** Human Genome Center, the Institute of Medical Science, the University of Tokyo, 4-6-1 Shirokanedai Minato-ku, Tokyo 108-8639, Japan

**Keywords:** genome organization, regression modeling, graph embedding, B cell differentiation, lymphoma

## Abstract

Delineating the intricate interplay between promoter-proximal and -distal regulators is crucial for understanding the function of transcriptional mediator complexes implicated in the regulation of gene expression. The aim of the present study was to develop a computational method for accurately modeling the spatial proximal and distal regulatory interactions. Our method combined regression-based models to identify key regulators through gene expression prediction and a graph-embedding approach to detect coregulated genes. This approach enabled a detailed investigation of the gene regulatory mechanisms underlying peripheral B cell differentiation, accompanied by dramatic rearrangements of the genome structure. We found that while the promoter-proximal elements were the principal regulators of gene expression, the distal regulators fine-tuned transcription. Moreover, our approach unveiled the presence of modular regulators, such as structural cofactors and proximal/distal transcriptional factors, which were co-expressed with their target genes. These findings imply that the dysregulation of interactions between transcriptional and structural factors is associated with chromatin reorganization failure and ultimately an increased risk of malignancy. We envisage that our computational approach will help crack the transcriptional *cis*-regulatory code of the three-dimensional network regulating gene expression.

## Introduction

Genes of multicellular organisms are regulated via a diverse set of mechanisms, which are often cell-type specific. Growing evidence supports the notion that the dynamic rearrangement of the spatial genome architecture affects the interactions among *cis*-regulatory elements in the nuclear microenvironment, constituting the so called ‘transcriptional domain’. Within this domain, distal regulatory elements (e.g., enhancers, silencers, and insulators) communicate with promoter-proximal regulatory elements and exhibit modularity, cooperativity, and specificity in spatiotemporal control of gene expression (1–3). Therefore, deciphering the *cis*-regulatory code in the three-dimensional (3D) transcriptional domain is fundamentally important for understanding the diverse roles and mechanisms of gene regulation.

Over the years, many studies have investigated the roles of 3D regulatory interactions in processes such as cell-fate decision and disease development (4,5). For instance, dramatic genome rearrangement occurs during the transition from naive B (NB) cells to germinal center B (GCB) cells. This rearrangement involves many locus control regions and enhancers, which effects the expression of thousands of cell type-specific genes (6–8). However, our understanding of the roles of enhancers and associated signaling transfer during promoter targeting is still very limited. One of the primary challenges is the sheer complexity of the gene regulatory networks (6,9), which are composed of gene regulators interacting with multiple transcriptional and structural proteins (5,10,11). Although recent computational approaches have made considerable breakthroughs in characterizing the genome sequences of important regulatory elements (12,13), there has been less focus on dissecting their multiway interactions.

The aim of the present study was therefore to use a computational method to model the interactions among promoter-distal and -proximal regulators and associated structural cofactors, collectively referred to as the ‘3D regulatory interaction’ of gene expression. After defining the 3D transcriptional domains using Hi-C (High-throughput Chromosome Conformation Capture) contact information, we used a regression-based algorithm to infer key factors implicated in gene expression regulation (14).

A graph-embedding algorithm (15), which characterized Cofactor-TF-Gene networks, was subsequently used to detect coregulated sets of cell type-specific genes and their regulators. By applying this novel approach to the investigation of human B cell response regulation, we discovered that the interplay between transcriptional and structural factors is essential for the function of 3D regulatory networks within the transcriptional domain. Our findings highlight the importance of understanding the dynamics of spatial regulatory interactions that could help improve the accuracy of disease mapping using modular biomarkers.

## Materials and Methods

### Data preparation

The raw and processed NB and GCB datasets were downloaded from the Gene Expression Omnibus (GEO), under accession numbers GSE84022 (for RNA-seq FASTQ files), GSE159314 (for ATAC-seq narrow peaks), and GSE84022 (for Hi-C contacts). Data relating to the narrow peaks of the six histone marks H3K4me1, H3K4me3, H3K9me3, H3K27me3, H3K36me3, and H3K27ac were downloaded from DeepBlue (16). The RNA-seq FASTQ files generated using the following cancer cell lines were obtained from CCLE (17): Ly10, activated B-cell-like (ABC) subtype of diffuse large B-cell lymphoma (DLBCL); Ly7, GCB-cell subtype of DLBCL; BL-41, Burkitt lymphoma; and Mino, mantle cell lymphoma. Data relating to the ChIP-seq peaks of *BACH2* (GSM1084800), *MAFK* (GSM1159670), and *MYC* (GSM762710) were also downloaded from GEO. The experimentally validated data relating to the physical protein-protein interactions (PPIs) were retrieved from STRING (18) with a confidence score (PPI score) > 0.4.

### Data processing

After assessing the RNA-seq reads using Trimmomatic (ver. 0.39) with the option “ILLUMINACLIP:adapter_file:2:30:10 LEADING:20 TRAILING:20 MINLEN:36” (19), HISAT2 (ver. 2.2.1) (20) was used to align the quality-controlled reads to the hg38 human reference genome. Cufflinks (ver. 2.2.1), with RefSeq coding-gene annotation (21), was used to quantify gene expression levels in terms of fragments per kilobase of exon per million reads mapped (FPKM) by merging multiple replicates. All the FPKMs were then normalized to transcripts per million (TPM) values. The peak positions of the ChIP-seq and ATAC-seq datasets were retrieved using q-value < 0.05 and a > 2 fold-change (FC); regions that did not overlap with the blacklisted genomic regions were retained (22).

The contact frequencies of intra-chromosomal Hi-C anchor pairs were normalized by the Vanilla-Coverage approach (23). When the center of an anchor located at intergenic regions connected with its partner anchor situated more than 10 Kb away and at a core promoter, spanning −2,500 bp to +500 bp from a transcription start site (TSS), the center of that intergenic anchor was bidirectionally stretched to a 5-Kb window and defined as a long-range contact (LRC) for a corresponding gene.

### Scoring transcription factor binding sites (TFBSs) in the promoter region

MATCH in the minimize false-positive mode (24) with TRANSFAC DB (https://www.genexplain.jp/transfac.html) was used to scan for TFBSs from the DNA sequences of the proximal promoter region (−10 Kb from the TSS). Considering that a TFBS may be detected repeatedly from a promoter region and bound by multiple transcription factors (TFs) during gene expression, a TFBS *j* which was found *K* times within the promoter *i* and potentially targeted by *N* number of TFs was scored as follows:

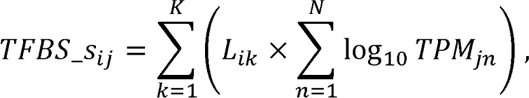

where *L_ik_* represents a weight based on the location of the *k*th TFBS*_j_* in the promoter *i*, and TPM*_jn_*is defined using histograms displaying,_obs_, which represents the distribution of TFBS*_j_* positions the expression level of TF*_n_* that binds to the TFBS. As in previous studies (14,25), the weight *L_ik_* was observed across all the promoters, and,_rnd_, which represents the random distribution of TFBS*_j_*, calculated as follows:

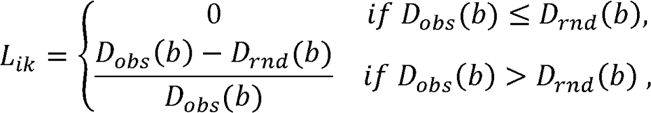

where *b* corresponds to the index of 100-bp bins in which the location of the *k*th TFBS*j* is contained.

### Scoring TFBSs in the LRC region

Considering that a promoter may contact multiple 5-Kb LRCs, TFBSs found within LRCs (i.e., lrcTFBSs) may appear multiple times within the same or originate from different LRCs. Thus, a lrcTFBS *j* found *K* times from the *C* number of LRCs contacting the promoter *i* was scored as follows:

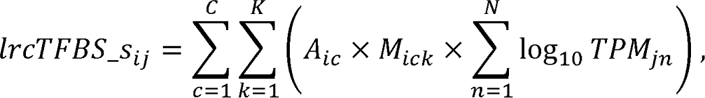

where *A_ic_*stands for the activity of the *c*th LRC for the promoter *i*, and TPM*_jn_*is the expression level of TF*_n_* binding to the TFBS. *M_ick_* represents the MATCH score (24), which is a score, ranging from 0.7 to 1.0, used to grade the alignment of lrcTFBS*j* with the LRC sequences. As in the Active-by-Contact model (11), the activity *A_ic_* was calculated as:

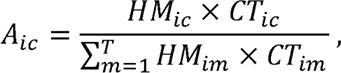

where *T* represents the total number of distinct LRCs contacting the promoter *i*, and *HM* and *CT* stand for the fold-enrichment (FE) value of histone modification and the normalized Hi-C contacts, respectively. The histone FE is the mean of the log-2 FE signals of H3K27ac and H3K4me1 peaks found within the 5-Kb LRC regions.

### Building regression models

Given a set of TPM values as response variables, a linear regression model predicts the TPM value with *n* number of explanatory variables (i.e., regulatory features) by estimating regression coefficients (RCs) as follows:

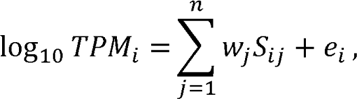

where S*_ij_* _and_ w*_j_* represent a regulatory feature *j* for the gene *i* and a regression coefficient for the feature *j*, respectively. Conducting comparative studies using previously published models is challenging and may lead to biased interpretations. To address this issue, new models were constructed specifically in the present study. These models used four distinct sets of regulatory features, each of which had a different number of explanatory variables: 1) the baseline model (B model) using *TFBS_s_ij_* within 10-Kb TSS-upstream regions; 2) the BH model was an extended version of the B model, which also included the log-2 FEs of each of the six histone markers located in the promoter regions; 3) the BL model was an extended version of the B model, which was generated by additionally incorporating *lrcTFBS_s_ij_*, and 4) the BHL model was constructed by combining the BL and BH models. Randomized versions of the above models were also generated.

### Feature selection procedure

An iterative procedure was used to find statistically significant explanatory variables through a greedy search (Figure S1) as previously described (14). In brief, stepwise model selection, based on Akaike’s Information Criterion (AIC), was used to predict the most adequate explanatory variables from the full regression model. This process yielded the reduced model M1, the predictive power of which was evaluated using 5-fold cross-validation (CV). Next, the M1 was extended by introducing a variable randomly selected from the set of variables that AIC had removed in the previous step; this process generated the trial model M2. The performance of M2 was assessed using AIC and the 5-fold CV. If M2 demonstrated improved predictive power over M1, it was selected; otherwise, another M2, with a different random variable, was subsequently tested. This procedure was repeated until all the variables removed during the AIC step had been tested. Each iteration of this feature selection procedure was performed 10 times by specifying different random seeds, which yields an ensemble of RCs.

### Graph embedding

Using significant TFBSs identified by the regression model, a network of TF-Gene interactions was constructed and subsequently extended to form a network of Cofactor-TF-Gene interactions. The inclusion of TFs and cofactors in this analysis was contingent on the expression of their coding genes in NB and/or GCB cells at levels > 3 TPM. The cofactors had to physically interact with at least one TF. According to the graph-embedding method, specifically the Large-scale Information Network Embedding (LINE) model (15,26), each vertex in the network was represented by a 200-dimensional vector based on the similarity of sub-graph structures. For instance, in a binary-weighted undirected graph, the one-step neighbor of a vertex (i.e., the first-order proximity) and the neighbor’s neighbor of the vertex (i.e., the second-order proximity) are preserved in a low-dimensional dense vector space. To optimize the embedding vectors, an asynchronous stochastic gradient algorithm with negative edge sampling was used to minimize the two objective functions of the first-(*O_1_*) and second-order (*O_2_*) proximities, based on KL-divergence as a distance function:

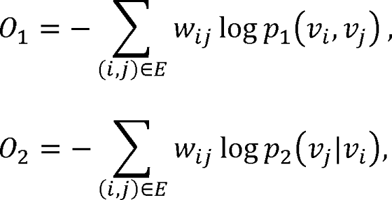

where *E* represents the edge set, *p_1_* is the joint probability between vertex *v_i_* and *v_j_*, and *p_2_* is the conditional distribution. The LINE package (https://github.com/tangjianpku/LINE) was used to reconstruct the input network with a parameter “-depth 2”. LINE models were trained for each of the orders with the default parameter “-size 100 -negative 5 -rho 0.025”, before model normalization and concatenation were performed. The dimensionality of the embedding vectors was reduced using the t-SNE algorithm, before performing K-means clustering with the t-SNE vectors.

### Bioinformatics analysis

All the publicly available datasets were converted into the hg38 coordinate using UCSC LiftOver (ver. 3.0). After removing the human housekeeping genes defined at HRT Atlas (27), EdgeR (28) was used to detect the differentially expressed genes (DEGs), including differentially upregulated genes (DUGs) and differentially downregulated genes (DDGs), which satisfied the > 2 FC and < 0.05 false discovery rate (FDR) criteria. The FDR was estimated using the Benjamini-Hochberg correction. Two-tailed t-tests were used to estimate the significance of the ensemble of regression coefficients. After applying Bonferroni correction, explanatory variables with an adjusted p-value < 0.05 were considered to be statistically significant.

BEDtools (ver. 2.25) (29) was used to manipulate peaks, including intersecting and extending them within the genomic context. Cytoscape (ver. 3.9.1) (30) was employed to visualize the networks. The residual standard error refers for the square root of an overall difference between TPM values and predictions, which was calculated by dividing the residual sum of squares (RSS) by the degree of freedom (i.e., the total number of genes – 2). The following packages in R language (https://www.r-project.org/) were used to perform the analyses and data visualization: ‘enrichGO’ in the ‘clusterProfiler’ package with a parameter “pAdjustMethod=BH, pvalueCutoff=0.01, qvalueCutoff=0.05” for testing the enrichment of Gene Ontology (GO) biological process (BP) terms; ‘Rtsne’ with the default parameter along with principle component analysis (PCA) for dimensionality reduction; ‘lm’ and ‘stepAIC’ for fitting regression models; and ‘hclust’ with the average-linkage method in Euclidean distance for hierarchical clustering.

## Results

### Overview of transcriptional regulation modeling

To profile the promoter-proximal and -distal regulators of specifically expressed genes, we initially explored the intra-chromosomal interactions captured by Hi-C (6). We found that, in NB cells, 48.2% and 38.3% of the 133.2 million Hi-C contacts were located within the intergenic and gene-body (GB) regions, respectively, and were paired with promoters in the regions 10-Kb upstream of the TSS. Similarly, in GCB cells, 40.4% and 35.5% of the 54.5 million Hi-C contacts were promoter-intergenic and promoter-GB anchor pairs, respectively. By merging the 1-Kb intergenic Hi-C anchors that overlapped with each other and were located at > 10 Kb from their core promoter (−2,500 bp to +500 bp from the TSS), we were able to identify the paired anchors linking the intergenic regions to the core promoters. Thus, in NB cells, we identified 577,857 paired anchors contacting 7,918 intergenic regions and 11,678 promoters, while in GCB cells, we identified 415,395 paired anchors contacting 7,870 intergenic regions and 11,360 promoters.

Next, we increased the intergenic anchor window to within 5 Kb from the center and defined these regions as LRCs; LRCs shape the genomic regions closely positioned within the 3D transcriptional domain of a gene (Figure 1A). Lastly, we scanned the DNA sequences of promoters and LRCs for TFBSs and scored the TFBSs within the promoter (prmTFBSs) or LRC (lrcTFBSs) regions. As described in previous studies (14,25), the scoring function for prmTFBSs takes into account the non-random positioning and the expression levels of the TF genes that have the potential to bind to the prmTFBSs (Figure 1B). In addition, we scored the lrcTFBSs by estimating their LRC activities (Figure 1C). The lrcTFBS scoring method follows a similar principle to the activity-by-contact model (11), wherein an LRC is weighted according to the Hi-C contact frequency, along with the extent of H3K27ac and H3K4me1 enrichment (see the Methods section for details).

**Figure 1.**
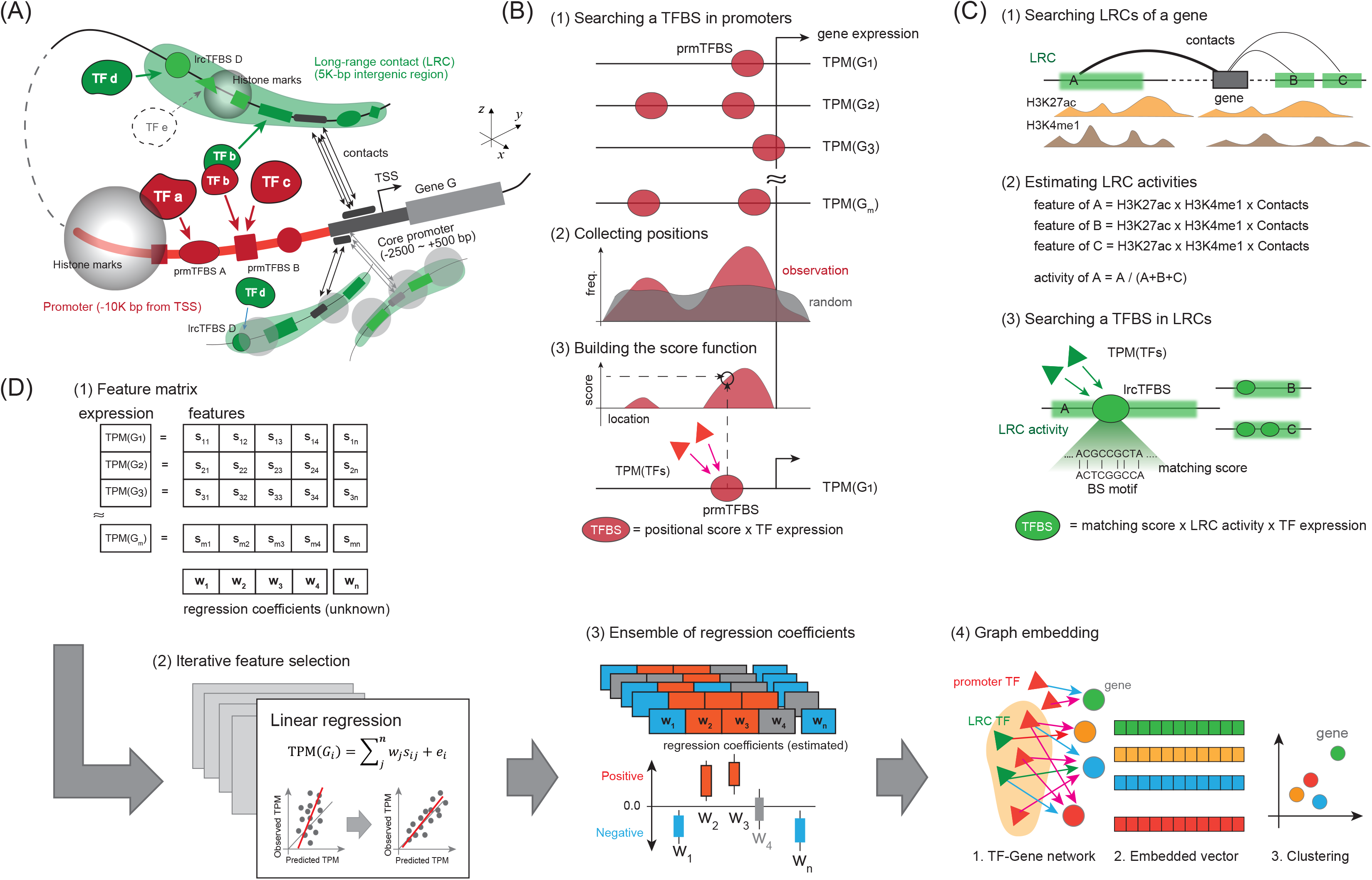
Overview of the computational pipeline to model gene expression regulation. (A) Schematic representation defining the 3D transcriptional domain for a gene to be modeled. (B) Scoring of proximal promoter TFBSs (prmTFBSs) by using positional information and the expression of TF genes. (C) Scoring of distal intergenic TFBSs (lrcTFBSs) based on the LRC activity and the expression of TF genes. (D) The analytic pipeline used to infer which regulators are important for gene expression using linear regression modeling and to identify coregulated genes using a graph-embedding method. LRC, long range contact; TF, transcription factor; TFBS, TF binding site; TPM, transcripts per million; TSS, transcription start site.

To identify critical regulatory elements and determine how they interact in the gene regulatory network, we built an analytic pipeline that integrated generalized linear regression modeling and graph embedding (Figure 1D). The pipeline followed a systematic approach referred to as greedy iterative feature selection, to identify the explanatory features essential for predicting gene expression (14). The feature selection process was iteratively repeated to produce a collection of regression coefficients (RCs) (Figure S1). After performing statistical tests with these RC ensembles, we constructed TF-Gene regulatory networks and mapped these networks to a low-dimension vector space using the LINE graph-embedding method (15,26). This approach enabled the clustering of genes that share regulatory interactions. Furthermore, we extended the TF-Gene networks by including cofactors, which represent tertiary regulatory complexes in the 3D transcriptional domain, as recently reviewed as an important factor in gene regulation (10,31,32).

### Genetic and epigenetic profiling in B-cell differentiation

We identified 1,749 DEGs that satisfied the > 2 FC and < 0.05 FDR criteria: 648 DUGs of NB cells and 1,101 DUGs of GCB cells (Figure 2A). The GO term enrichment analysis confirmed that the DUGs in GCB cells (GCB-DUGs) were involved in cell proliferation as reported previously (33) and also observed in lymphoma cells (Cluster 1 in Figure 2B). In addition, we found a subset of GCB-DUGs that was neither upregulated in lymphoma cells nor enriched in any of the GO terms (Cluster 3 in Figure 2B).

**Figure 2.**
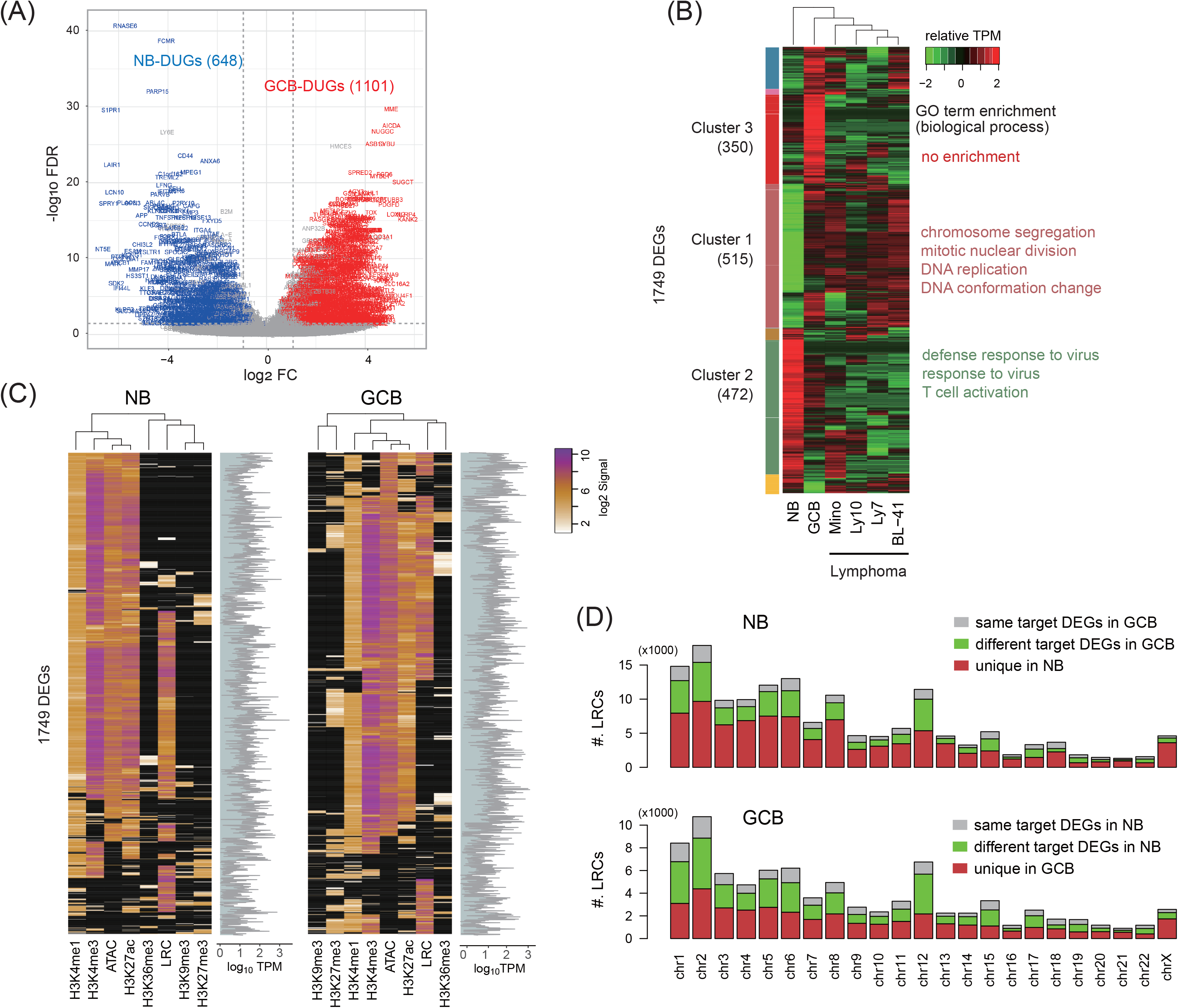
Genetic and epigenetic profiles in B cell sub-types. (A) Volcano plot showing DEGs between naive B (NB) and germinal center B (GCB) cells. (B) Clustering DEGs according to their expression patterns in healthy B cells and lymphoma cells. (C) Genetic and epigenetic profiles of DEG promoters along with the expression TPM values. (D) Distribution of LRCs targeting DEGs conserved or altered between the cell types. DEGs, differentially expressed genes; DUGs, differentially up-regulated genes; FC, fold change; FDR, false discovery rate; LRCs, long-range contacts; TPM, transcripts per million.

Next, we conducted epigenetic profiling, which included six different histone modifications, chromatin accessibility, and LRC frequency, around the core promoters of the DEGs (Figure 2C). The results indicated that these DEG promoters were mostly highly accessible and exhibited active histone signatures (i.e., H3K4me1, H3K4me3, and H3K27ac); however, none of these epigenetic signals clearly correlated with the gene expression levels. In addition, we found that although 60% of the DEG promoters had LRCs (Figure 2C), more than 80% of these LRCs were cell type-specific (Figure 2D). Specifically, 86.7% of the 115,840 NB-LRCs and 81.7% of the 71,472 GCB-LRCs were exclusively observed in or linked to different gene promoters (Table S1).

Collectively, our epigenetic profiles support the notion that peripheral B cell differentiation is accompanied by the cell type-specific reorganization of 3D genome structures and the cell cycle initiation (34,35). Furthermore, our findings imply that the intricate interplay between multimodal genetic and epigenetic factors is required for tight gene regulation.

### Identifying critical regulators by predicting gene expression

We next identified 78 lrcTFBSs and 77 prmTFBSs among the NB-DEGs, as well as 79 lrcTFBSs and 78 prmTFBSs among the GCB-DEGs. We built and tested eight regression models using different combinations of feature sets and target genes, in addition to the random shuffling of these features. A feature set containing only prmTFBSs was used as the baseline model (B model). The B model was expanded by using a set of prmTFBSs and histone enrichment markers in the promoter region to generate the BH model. The BL model was constructed using a set of prmTFBSs and lrcTFBSs. Finally, the BHL model combined the features of the BH and BL models.

The BH model, which considers only the promoter configuration, significantly improved the predictive performance of the B and BL models (Figure 3A). Interestingly, the BHL model, combining the lrcTFBS features not only further improved the identification of DUGs but also that of DEGs in general (Figure 3B, Figure S2A). On inspecting the selected features in each model, we found that the BHL model was based on the combinatorial effect of histone modifications and proximal/distal TFBSs (Figure S2B). For example, in the prediction of GCB-DEG expression, 42 TFBSs were detected as having significant RCs (< 0.05 adjusted p-value). The activity of a given TFBS was calculated by multiplying the mean RC by the total TPM value of binding-TF genes. This assessment revealed whether a TFBS positively or negatively impacted the prediction of gene expression (Figure S2C), indicating its role as an activator or a repressor, respectively.

**Figure 3.**
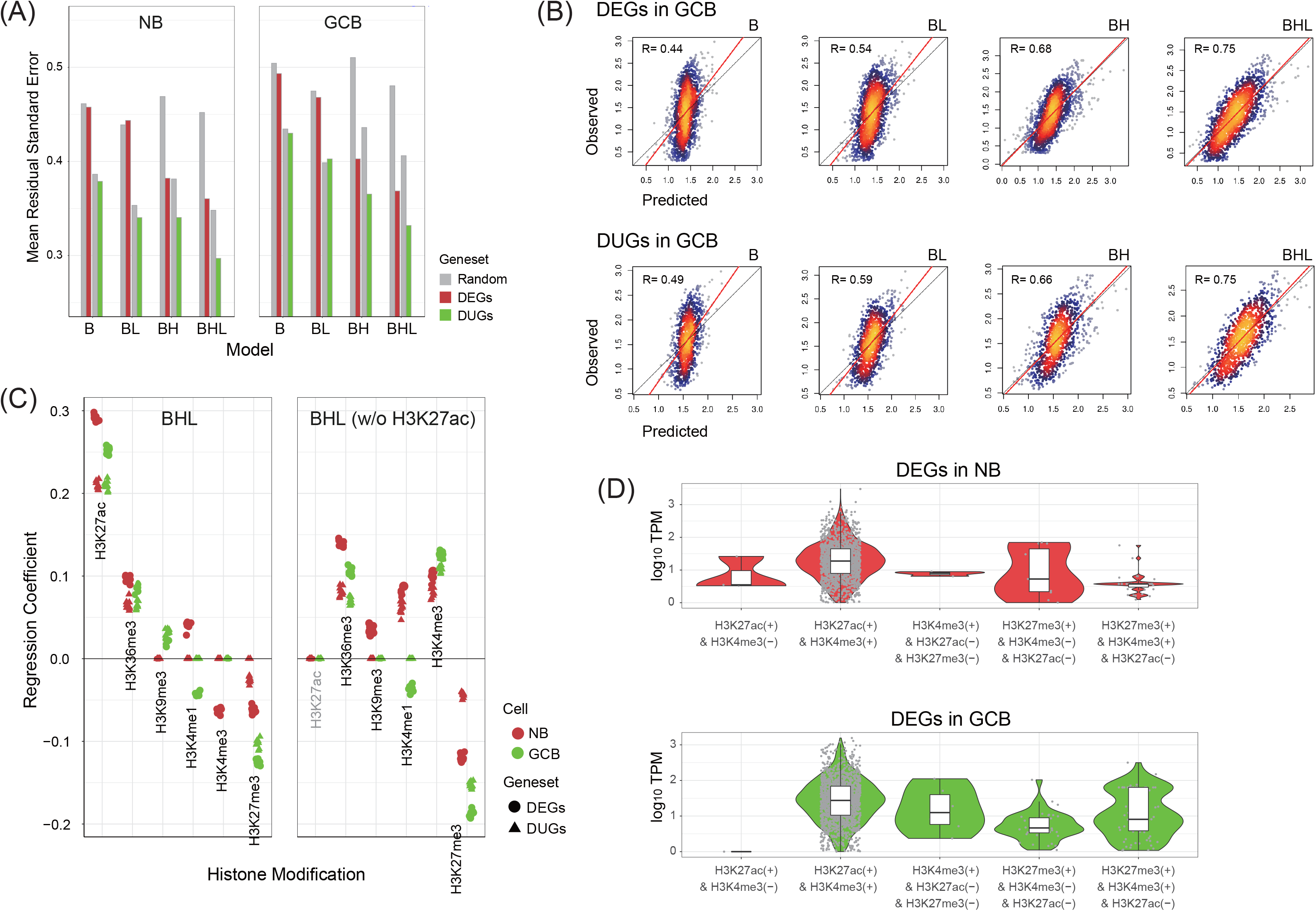
Performance of gene expression predictions and implication of histone modifications. (A) Comparing the average performance of the four models (B, BH, BL, BHL) in predicting the expression of different gene sets. (B) Scatter plots showing the correlations between the observed and predicted gene expression. (C) Distribution of the RC ensembles for the various histone marks with (left) and without (right) the inclusion of H3K27ac in the BHL model, which demonstrates the marked change in H3K4me3 RC. (D) Violin plots showing the combinatorial histone marks found on the DEG promoters, showing the dominant co-enrichment of H3K27ac with H3K4me3 and the bivalency of H3K27me3 and H3K4me3 for DEGs suppressed in NB versus GCB cells. B, the baseline model using prmTFBSs only; BH, the BH model using prmTFBSs and histone marks; BL, the BL model using prmTFBSs and lrcTFBSs; BHL, the BHL model combining the BH and BL models; R, Pearson’s correlation coefficient; RC, regression coefficient.

In the BHL model, the explanatory features H3K27ac and H3K27me3 were estimated with remarkable RCs (Figure S2B), which supports their importance in B cell differentiation (36). By contrast, H3K4me3, an active promoter marker, was associated with negligeable RCs or even negative RCs during the prediction of NB-DEGs (Figure 3C). To explain this observation, we next analyzed the distribution of histone markers in NB cells. We found that H3K27ac and H3K4me3 were frequently co-enriched during the expression of a wide range of genes (Figure 3D). This observation aligns with the transcriptional interplay previously reported between these two histone modifications (37,38). Of note, a subset of NB-DEGs, including those encoding cell adhesion molecules (e.g., protocadherin and LOXL2), were associated with the simultaneous presence of H3K4me3 and H3K27me3 but not H3K27ac, and their expression was particularly suppressed in NB cells. Therefore, it is reasonable to assume that the enrichment of H3K27ac was sufficient to explain the DEG expression, and the negative RCs of H3K4me3 likely accounted for the poised and repressed genes regulated by the bivalent chromatin domain (39,40). Indeed, removing H3K27ac from the model restored the significantly positive RCs of the H3K4me3 without sacrificing overall model performance (R=0.74) (Figure 3C).

Taken together, our results imply that a synergy exists between the actions of promoter-proximal and -distal regulatory elements within the 3D transcriptional domain, which are also influenced by histone configuration in B cell differentiation (41). Moreover, our data suggest that H3K27ac is required for the accurate modeling of poised gene regulation in NB cells.

### Inferring the modes of proximal and distal regulator effects

Our regression models successfully identified well-documented transcriptional *cis*-regulators, such as the TFBSs Bcl-6, AP-1, Blimp-1, Bach2, and XBP1 (6,42,43), along with the diverse TFBS activities, according to their locations and target genes (Figure 4A, Figure S2C). For instance, in the model predicting NB-DUG expression, Bcl-6 and Blimp-1 were identified in the promoter regions, while STAT1 and AP-1 originated from LRCs. Some TFBSs, such as REST, Homez, and MEF-2, were detected in both the promoters and LRCs, where they exhibited mutually opposing activities.

**Figure 4.**
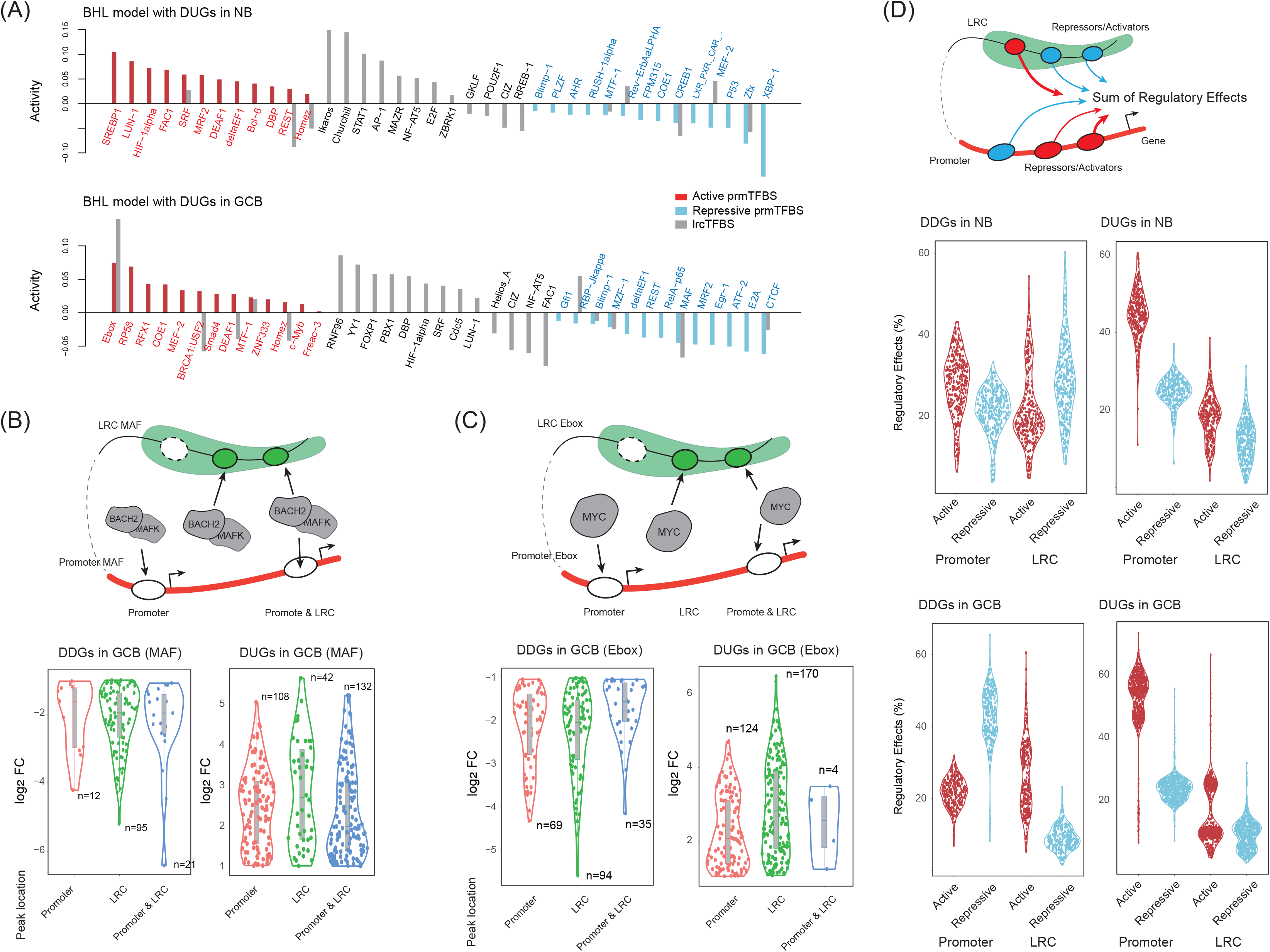
Contribution of proximal and distal regulatory effects inferred by regression models. (A) Distribution of the activities of prmTFBSs and lrcTFBSs in the prediction of DUG expression (< 0.05 adjusted p-value). (B) Schematic diagrams (top) showing the category of the presence of BACH2/MAFK binding peaks. Violin plots (bottom) showing the fold change in gene expression in GCB versus NB cells. (C) Schematic diagrams (top) showing the category of the presence of MYC binding peaks. Fold change (bottom) of gene expression in GCB versus NB cells. (D) Schematic diagram (top) and violin plots (middle and bottom) showing the proportions of active and repressive effects exerted by prmTFBSs and lrcTFBSs. DDGs, differentially downregulated genes; DUGs, differentially upregulated genes; FC, fold change.

To validate the inferred TFBS activities, we compared the publicly available peak positions of three TFs in GCB cells; *BACH2* and *MAFK* binding to the TFBS MAF (44), and *MYC* binding to the TFBS Ebox (45). Although the MAF was detected with negative RCs (Figure 4A), consistent with the repressive activities of *BACH2* and *MAFK* (42), most of the DDGs possessed the TF peaks at LRCs (Figure 4B). Interestingly, simultaneous peak presence at the promoters and LRCs was associated with gene suppression. For the *MYC* bindings, our model inferred the Ebox with positive RCs, consistent with its activating function (46). Unlike the MAF demonstrating the cooperative effect of promoter-bound and LRC-bound *BACH2* and *MAFK*, most of the *MYC* peaks were positioned at either the promoter or LRC regions (Figure 4C).

Next, we analyzed the TFBS activities by categorizing their locations and their modes of activation or repression (Figure 4D). Overall, both NB-DUGs and GCB-DUGs primarily received positive regulatory effects originating from their promoters, which suggests the primary contribution of promoter-proximal regulators in gene upregulation. Conversely, the NB-DDGs were predominantly repressed by LRC-mediated signals, which only affected a small portion of GCB-DDGs. This finding may indicate the presence of a poised but repressed 3D genomic configuration in NB cells, which subsequently undergoes activation in GCB cells (7).

Collectively, these results show that 3D genome reorganization regulates the dynamic function of proximal and distal *cis*-regulatory elements during the NB to GCB transition. Moreover, the prmTFBSs appear to be the main regulatory effectors and the lrcTFBSs fine-tune transcription, consistent with a previous study (47).

### Characterizing 3D regulatory modules

To explore how the 3D transcriptional domain combines the activities of promoter-proximal and - distal TFs, we constructed gene regulatory networks comprising TFs with prmTFBS and/or lrcTFBS binding potential (named and prmTFs and lrcTFs, respectively). We identified 147 TFs with > 3 TPM, of which 38% were DEGs, comprising 82 prmTFs and 65 lrcTFs. Because multiple TFs can regulate a single gene, the resulting network was highly complex (Figure 5A). Thus, we employed a graph-embedding method (26) to convert the nodes (i.e., prmTFs, lrcTFs, and DUGs) into 200-dimensional vectors based on the sub-graph structures; these vectors were then clustered using a K-means algorithm as previously described (15). This approach clearly separated the nodes between the cell types and within a cell type, which implies the presence of distinctive TF-Gene connectivities (Figure 5B).

**Figure 5.**
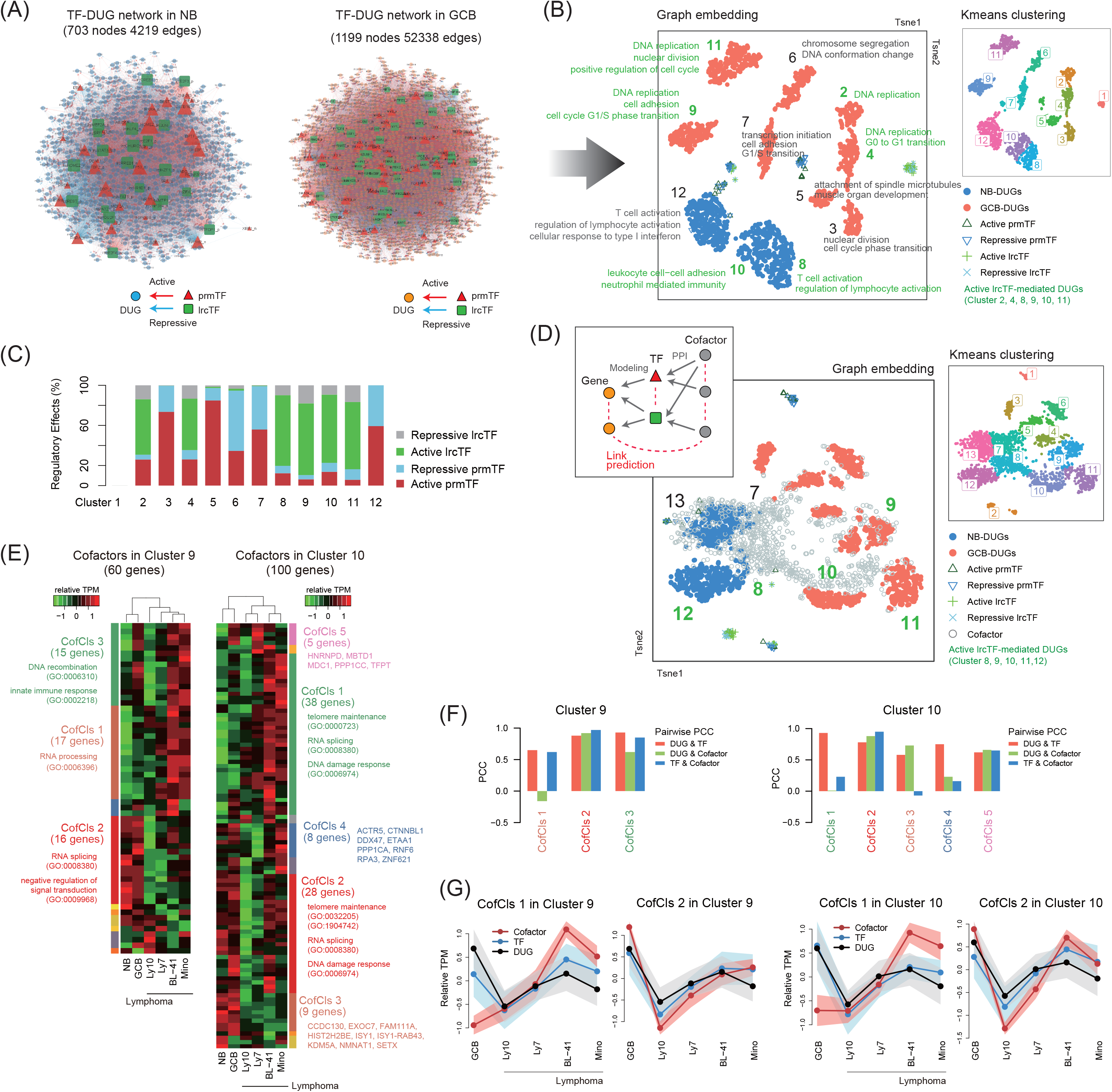
Results of the graph-embedding method with gene regulatory networks and the expression levels of clustered genes in lymphoma. (A) TF-Gene regulatory networks inferred by the regression modeling. (B) Two-dimensional t-SNE plots depicting the embedded nodes of the TF-Gene networks. The DEGs in each cluster share TF-Gene connectivity, which implies co-regulation, and are enriched in GO terms (< 0.05 q-value), highlighted in green for lrcTF-involved clusters. (C) Proportion of activating and repressive effects exerted by the prmTFs and lrcTFs in each DEG cluster shown in (B). (D) Two-dimensional t-SNE plots depicting the embedded nodes of the Cofactor-TF-Gene networks. (E) Hierarchical clustering of cofactor expression in healthy B cells and lymphoma cells. The cofactors were derived from the corresponding clusters shown in (D). (F) Correlations among the mean expression levels of Cofactors, TFs, and DUGs from Clusters 9 and 10 of (D) in GCB and lymphoma cells. (G) Correlated expression patterns exemplifying the Cofactor-TF modules that regulate the target GCB-DUGs. CofCls, cofactor cluster; TF, transcription factor; PCC, Pearson’s clustering coefficient; PPI, protein-protein interaction.

Next, we calculated TF activity scores by multiplying the mean RC of the TFBS by the TPM of the TF-coding gene. We found the markedly positive regulatory effects of lrcTFs in Clusters 2, 4, 8, 9, 10, and 11 (Figure 5C). Of note, although the 350 GCB-DUGs of Cluster 3, shown in Figure 2B clustering genes based on the expression profile, were not enriched in any GO terms, these genes were scattered across all the GCB-related clusters and enriched by crucial biological processes in Figure 5B (Table S2). These results imply that the TF-Gene regulatory connectivity model is better at classifying functionally relevant genes than expression-based models. Interestingly, a specific GO term was repeatedly enriched in the different clusters as shown in Figure 5B. This result suggests that multiple sets of distinctive TF-Gene interactions regulate the biological processes in B cell differentiation.

Given that several studies have reported the functional importance of cofactors in enhancer-promoter communication (48,49), we extended the TF-Gene networks by adding 852 cofactor genes expressed in the cells (> 3 TPM). These genes were non-DEGs and annotated in STRING (18) as physically interacting with prmTFs and/or lrcTFs (> 0.4 PPI score). After embedding and clustering the nodes, we identified DUGs and co-clustered cofactors (Figure 5D). For example, Clusters 8 and 12 comprised NB-DUGs, while Clusters 9, 10, and 11 included GCB-DUGs. These clusters also contained unique cofactors that interacted with DUG-targeting TFs (Figure S3A) and were involved in the essential biological processes in B cells (Figure S3B).

Unexpectedly, the expression of cofactors was markedly altered in lymphoma versus normal B cells, and these changes were associated with the clustering patterns observed in these cells (Figure 5E). Moreover, some of the cofactor clusters (CofCls) correlated with the expression of their corresponding TFs and even DUGs targeted by these TFs (Figure S3C-D). For example, the genes within CofCls 2 in Clusters 9 and 10, which were associated with RNA processing and the maintenance of chromosome structure, were expressed in conjunction with their interacting genes (Figure 5F). This finding highlights the important role of cofactors, which are coregulated with proximal/distal TFs to form a 3D regulatory network. Interestingly, although CofCls 1 in Clusters 9 and 10 was associated with similar biological functions as CofCls 2 and repressed in normal B cells, the expression of cofactor genes in CofCls 1 exhibited correlations with the interacting genes in lymphoma cells (Figure 5G). Therefore, in line with the findings of a previous study (10), it is plausible that the dysregulation of cofactors disrupts cofactor-TF interactions in the 3D transcriptional domain and contributes to cancer development.

### Identifying the regulatory modules for BCL6

To investigate the cofactor-TF modularity in more detail, we next focused specifically on *BCL6*, which is a master TF of GCB cells known to be controlled by LRCs (6,8). Overall, the regulatory network of *BCL6*, a GCB-DUG, comprised 13 prmTFs and 44 lrcTFs interacting with 681 cofactors (Figure S4A). Among these regulators, the graph-embedding approach clustered *BCL6* and 99 cofactors, which belonged to Cluster 10 (Figure 5D) and interacted with 16 TFs (Figure 6A). To confirm the presence of *BCL6* neighboring genes in the embedding space, we calculated the Pearson’s correlation coefficients (PCC) for the embedded *BCL6* vector and that of each of the 1,748 DEGs (Table S3). As expected, we found that the higher PCCs were derived from the DEGs in Cluster 10 (Figure 6B). Specifically, we identified 86 GCB-DUGs which exhibited an exceptionally high degree of similarity in their embedded vector representations (PCC > 0.99). This implies that the cofactors and TFs targeting *BCL6* (Figure 6A) coregulated these 86 GCB-DUGs, including cadherins and various enzymes.

**Figure 6.**
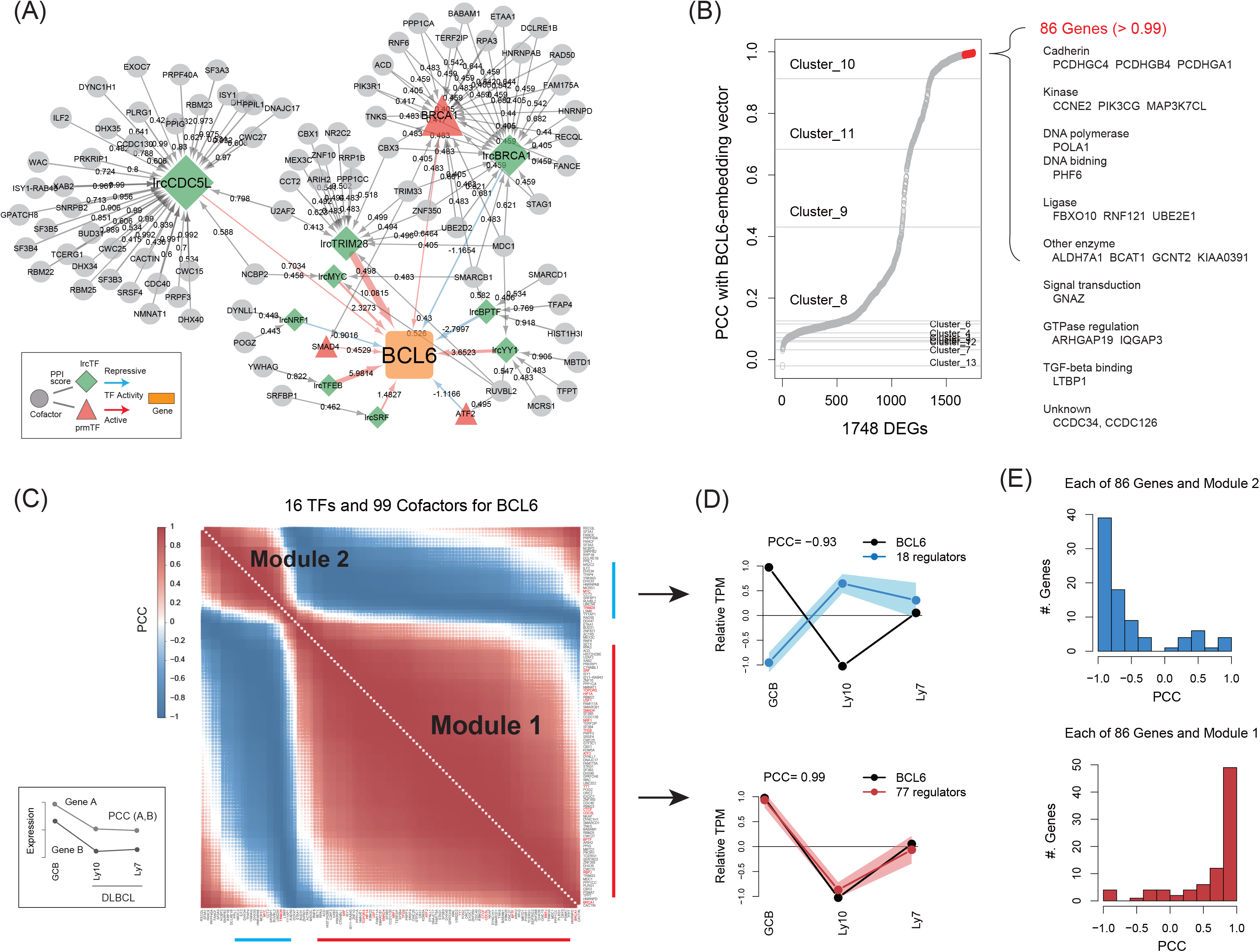
Cofactor-TF gene sets regulating BCL6 and presenting modular expression patterns in lymphoma. (A) The Cofactor-TF connectivity network characterized Cluster 10 of Figure 5D and was inferred to regulate *BCL6*. (B) Distribution of correlation coefficients between the graph-embedding vector of *BCL6* and that of each of the DEGs, based on the graph structure presented in Figure S4A. The cluster IDs correspond to those in Figure 5D. (C) Two regulatory modules of (A) showing similar expression pattern trends in GCB and lymphoma cells. (D) Correlation between the expression of *BCL6* and that of each of the modulated genes (> 0.5 and < −0.5 PCCs) in GCB and DLBCL cells. (E) Distribution of gene expression correlation coefficients between *BCL6* and each of the 86 coregulated genes shown in (B) in GCB and DLBCL cells. DLBCL, diffuse large B cell lymphoma; PCC, Pearson’s correlation coefficient.

The expression patterns of the 16 TFs and 99 cofactors (Figure 6A) in lymphoma cells frequently correlated with each other to form two regulatory modules (Figure 6C, Figure S4B-D). In DLBCL, where *BCL6* serves as a therapeutic target (8), 77 regulators of the major module underwent a reduction in expression in a manner similar to *BCL6*; meanwhile, 18 regulators of the other module exhibited the opposite expression pattern (Figure 6D). These correlations were also observed for the 86 GCB-DUGs potentially coregulated with *BCL6* (Figure 6E). Collectively, our findings suggest that the cofactor-TF modules tightly regulate target genes as activators or repressors. Thus, the abnormal switching of those modules could potentially increase the risk of malignancy, particularly when accompanied by dysregulated chromatin reorganization due to change in the expression of chromatin remodelers, such as *YY1*, *CTCF*, *STAG1*, *TRIM28*, *TFAP4*, and *RAD50* (Table S4).

## Discussion

To gain insights into the intricate connections between promoter-proximal and -distal regulators, we modeled the interactions among cofactors, proximal/distal TFs, and cell type-specific genes, which were associated with spatial genome arrangement in peripheral B cells during differentiation. We initially identified the TFBSs with the potential to bind TFs expressed in the 3D transcriptional domain, which were defined via cell type-specific Hi-C contacts. Next, we profiled the TFBSs as non-redundant prmTFBSs and lrcTFBSs by scoring their characteristics. Our regression models subsequently used these data to recapitulate the gene expression profiles and successfully quantify the parameters associated with their combinatorial regulation. Moreover, we constructed highly complex regulatory networks and used them to detect the specific sub-networks shared by functionally relevant gene subsets.

We next identified the overarching trends through a comprehensive analysis of our computational modeling data. For instance, the predominantly activating effect of promoter-proximal elements upregulated 1,749 DEGs in the B cells. The crucial function of distal elements, which further fine-tune transcription in a context-dependent manner, was subsequently exemplified by the MAF and Ebox TFBSs. Notably, the spatial genome arrangement, associated with specific cell states (e.g., the quiescent NB cell state), was likely responsible for the distal repressive effects observed. In total, 1,101 NB-DDGs were modeled in association with LRC-specific repressors in NB cells. These genes were upregulated during the differentiation of NB cells into GCB cells, in a process which was accompanied by the maintenance of pervasive H3K27ac-enriched chromatin, as well as a reduction in bivalent histone modifications and LRC-mediated repression. These findings underscore the importance of activators and repressors encoded in LRC regions (50), which are indispensable for the precise control of cell differentiation by interacting with B cell-lineage-specific epigenetic processes (2,51).

At the individual gene level, our results demonstrate that the complex 3D regulatory network, consisting of proximal/distal TFs and cofactors, exhibits modularity, which is essential for regulating the expression of target genes. For example, *BCL6* expression in GCB cells was modeled using 58 TFs interacting with 681 cofactors, which formed a unique 3D regulatory module comprising 16 TFs interacting with 99 cofactors; this module was responsible for regulating a subset of GCB-DUGs. Remarkably, differentially coregulated DEG subsets exhibited similar functions, even though the specific connectivity characteristics of their 3D regulatory modules varied. This finding suggests that parallel pathways activated by distinct transcriptional protein complexes determine the level of cell type-specific gene expression required for the regulation of B cell identity. However, this functional redundancy is thought to increase the risk of cancer development (52). Indeed, we found that members of the 3D regulatory modules were frequently co-expressed with their target genes in lymphoma cells, although the causality of this relationship is still unclear. Nevertheless, understanding how changes in 3D regulatory networks contribute to disease will help the development of potential modular biomarkers.

## Conclusion

Our computational framework inferring the proximal and distal regulatory interactions with incorporating PPIs provides an alternative approach for deciphering transcriptional *cis*-regulatory code mediated by the 3D chromatin structures. Further systematic investigations of factors, such as promoter-promoter interactions, enhancer-promoter proximity, and inter-chromosomal interactions, are warranted to gain a deeper understanding of how the interplay between cell-specific *cis*-regulatory elements and the higher-order genome structure regulates gene expression.

## Declarations

### Ethics approval and consent to participate

Not applicable.

### Consent for publication

Not applicable.

### Availability of data and material

All data used in this study were downloaded from public databases. The processed results are available as Supplementary data. The code is available on GitHub (https://github.com/Park-Sung-

Joon/GLMGE).

### Authors’ contributions

SJP designed the experiments and performed data analysis. SJP and KN drafted the manuscript. Both authors read and approved the final manuscript.

## Acknowledgements

Computational resource was supported by Human Genome Center (The Institute of Medical Science, The University of Tokyo, Tokyo, Japan).

## Additional files

### Supplementary Figures

This file includes; the schematic representation of the greedy feature selection method through the iterative linear regression (Figure S1), the predictive performance of the regression modeling of gene expression in NB cells and the distribution of the estimated RCs and TFBS activities (Figure S2), the further analyses of embedded clusters shown in Figure 5D (Figure S3), the Cofactor-TF-BCL6 network inferred from the regression model and the distribution of gene expression correlation coefficients of the *BCL6* regulators in GCB cells and four types of lymphoma cells (Figure S4).

### Supplementary Tables

This file includes; the detailed information of cell type-specific LRC-gene pairs (Table S1), the distribution of the 350 GCB-DUGs (Cluster 3 in Figure 2B) within the graph-embedding clusters shown in Figure 5B (Table S2), the Pearson’s correlation coefficients for the comparison of the embedded vector of *BCL6* and that of each of the 1,748 DEGs (Table S3), the Pearson’s correlation coefficients for the comparison of the expression of *BCL6* and that of each of the modulated genes in GCB and DLBCL cells (Table S4).

